# Circadian clock in the lateral habenula affects motor function in mice through the regulation of daily rhythms in the nigrostriatal pathway

**DOI:** 10.1101/2024.08.17.608401

**Authors:** Cassandra Goldfarb, Nadav Baharav, Heng Jiang, Amanda Szubinski, Shimon Amir, Konrad Schöttner

## Abstract

Alterations to the dopaminergic system can have several consequences on behaviour such as the development of disorders like addiction, schizophrenia, and Parkinsons. The lateral habenula (LHb) is uniquely positioned to act on a large group of dopaminergic (DA) neurons, sending inhibitory signals both directly and indirectly to the ventral tegmental area (VTA) and the substantia nigra (SN). Additionally, the LHb houses a circadian clock that appears to function independently from the central circadian pacemaker. We investigated the role of the LHb as a pacemaker for the production and release of DA along the nigrostriatal pathway. Using male and female *Bmal1* floxed mice, we injected AAV-2/9 Cre-eGFP virus into the LHb to selectively knockout *Bmal1*, a clock gene essential for clock functioning. We found a significant impact on motor functioning in both male and female knockout mice. Analysis of daily rhythms of expression of circadian clock genes and genes involved in DA synthesis, and liquid chromatography coupled mass spectrometry for striatal DA measurements revealed blunting of rhythms in the dorsal striatum (DS) and (SN) of knockout animals, that may contribute to the observed behavioural phenotype. As proper functioning of the striatum is likely maintained by a mutual interaction of the circadian clock and DAergic system, these findings support that disrupting the LHb clock can impact functioning of the nigrostriatal DA pathway.

## INTRODUCTION

Mammals exhibit daily rhythms in physiology and behaviour generated by an internal timekeeping system comprising an interconnected network of biological clocks in the brain body. The system provides a temporal order of biological processes according to the time of day, allowing organisms to anticipate challenges imposed by the cyclic changes in the environment (de Assis & Oster, 2021).

Tissue-specific clocks generate daily rhythms through the expression and reciprocal interaction of circadian clock genes and their protein products (King & Takahashi, 2000). Because these oscillations vary in phase, period, and stability, they must be constantly reset to remain aligned with rhythms of other internal processes and the external environment. Through mutual interactions of central and peripheral clocks within an organism, the circadian system provides the temporal integrity of metabolic, endocrine, and behavioural functions needed to ensure systemic homeostasis in a rapidly changing environment, which is critical health and well-being (Dibner et al., 2009).

It is proposed that diurnal changes in neuronal networks and signalling pathways within the brain, including the synthesis and release of neurotransmitters such as dopamine (DA), are crucial for the regulation of behavioural processes. Studies on the dorsal striatum (DS), the major input structure of the basal ganglia, revealed a clear link between circadian dysregulation and altered behavioural phenotypes in animal models (de Zavalia et al., 2021; Landgraf et al., 2016a; Ozburn et al., 2017; Schoettner et al., 2022). This view has been supported by studies on human brain tissue from donors with diagnosed mental illness (Ketchesin et al., 2023). Despite the significance of diurnal rhythms in striatal function, however, relatively little is known about how these oscillations are sustained, given that nuclei of the basal ganglia are not intrinsically rhythmic (Landgraf et al., 2016a) and lack direct projections from the SCN (Dibner et al., 2009; Iijima et al., 2002). It has therefore been proposed that temporal cues must be relayed through autonomous or semi-autonomous oscillators located in separate brain areas (Becker-Krail et al., 2022a; Pradel et al., 2022a).

Among several extra-SCN oscillators that have been identified in the brain, the lateral habenula (LHb) has emerged as a promising target involved in the regulation of daily rhythms in the basal ganglia. Neurons of the LHb display intrinsic daily rhythms in clock gene expression and electrophysiological activity through input from hypothalamic regions, including the SCN and the retina (Salaberry et al., 2019; Zhang et al., 2016). Moreover, the LHb exerts direct and indirect control over the ventral tegmental area (VTA) and substantia nigra (SN), thus affecting the rhythms of DAergic circuits.

Recent studies have highlighted the role of DA in the regulation of diurnal rhythms in the basal ganglia (Brami-Cherrier et al., 2020; Hood et al., 2010; Imbesi et al., 2009; Pradel et al., 2022a; Sahar et al., 2010; Uz et al., 2005). Levels of extracellular DA in the striatum vary across the 24-hour day (Ferris et al., 2014; Paulson & Robinson, 1995; Smith & Kieval, 2000) as a result of daily changes in DA synthesis and release. The expression of tyrosine hydroxylase (TH), the rate-limiting enzyme in DA production, varies across the day (Chung et al., 2014), just like levels of monoamine oxidase A (Maoa) and DA transporters (DAT) in the striatum (Ferris et al., 2014; Hampp et al., 2008). Notably, daily oscillations in DA levels affect rhythms in clock protein expression in medium spiny neurons (MSNs) of the DS (Hood et al., 2010), suggesting that diurnal changes in DA tone within the nigrostriatal pathway serve as a timing cue to synchronize and maintain striatal rhythms.

We therefore conceptualize a pacemaking function of the LHb that drives diurnal oscillations of DA levels in the nigrostriatal pathway, thereby affecting clock function in downstream brain regions, such as the DS. We tested this hypothesis by a series of experiments utilizing a mouse model with a conditional knockout of *Bmal1* within the LHb. Our work demonstrates that mice with a LHb specific deletion of *Bmal1* display blunted rhythms of striatal DA, which is likely caused by a dysregulation of daily rhythms of DA synthesis in the SN. Strikingly, these alterations are associated with attenuated motor functioning in male and female mice. Therefore, this work demonstrates the critical role of the LHb circadian clock in the coordination of DAergic functions in the basal ganglia and the regulation of behaviour.

## RESULTS

### *Bmal1* ablation from the LHb reduces wheel running activity but does not influence central clock function and affective behaviours

Bilateral intracerebral infusion of Cre-expression viral vector led to a successful ablation of *Bmal1* from the LHb. Immunofluorescence imaging confirmed the absence of BMAL1 protein from GFP labelled cells of the LHb exclusively (Fig. 1A). Western blot analysis furthermore demonstrated the absence of BMAL1 from tissue of the LHb collected from animals infused with virus expressing Cre protein when compared to controls (Fig. 1C).

**Figure 1.**
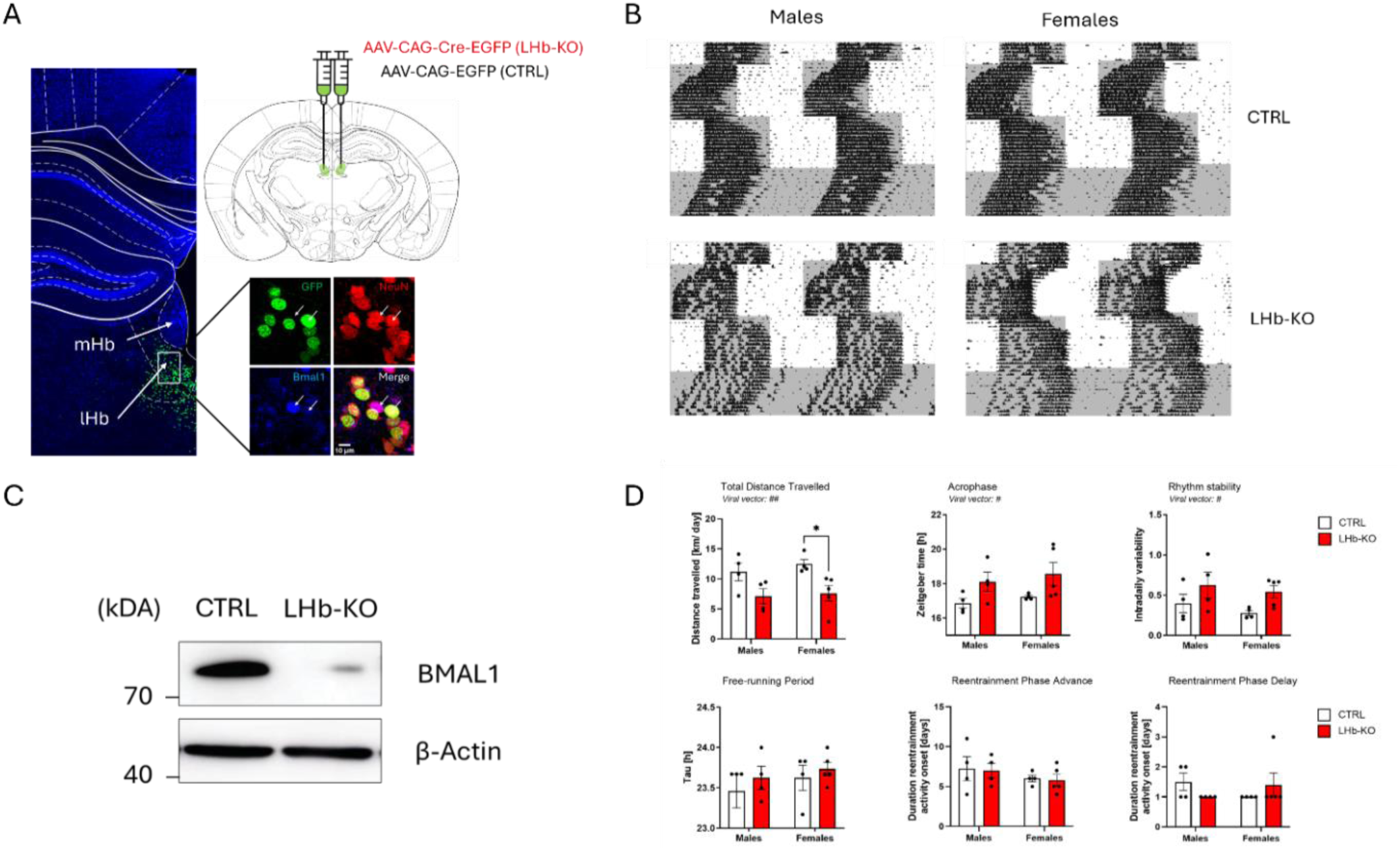
Bmal1 was successfully knocked out in the LHb. A) Representative immunofluorescence staining image of GFP expression in LHb of mice receiving stereotaxic injections of AAV-CAG-Cre-EGFP viral vector. 60x representative immunofluorescence staining in LHb tissue of a knockout mouse GFP (green), NeuN (red), and BMAL1 (blue) immunofluorescence staining in LHb tissue of a knockout mouse. Arrows demonstrate no overlap between GDP and BMAL1 in neurons. B) Representative actograms of mice under 12:12 LD after a 6- hour phase advance, 6-hour phase delay, and under constant darkness. C) Western blotting results levels of BMAL1 in pooled LHb tissue of KO and Ctrls, n = 4 per group. D) Running wheel data (males: n=8 (4KO); females: n=9 (5 KO)) was analyzed using Clocklab software, graph images depicting two-way ANOVA results for viral vector x sex: average daily distance travelled (*p< .05), acrophase, rhythm stability, free-running period, and time to entrain after a phase advance and phase delay are depicted.

To verify that the deletion of *Bmal1* did not affect central clock function, animals were kept in cages equipped with running wheels to study circadian rhythms of locomotor activity under various light/dark conditions. While free-running periods and re-entrainment durations following 6-h phase advances and 6-h phase delays of the light/dark cycle were unaffected by the deletion of *Bmal1* from the LHb (Fig. 1D, lower left, mid and right panels, Table. 1), levels of wheel running were reduced in knockout animals (Fig. 1D, upper left panel, Table. 1).

**Table 1.** Results of statistical analysis of wheel running activity and motor function in CTRL and KO mice.

| Locomotor activity rhythm |  | Two-way ANOVA |  |
| --- | --- | --- | --- |
|  | Sex | Viral vector | Interaction |
| Total distance travelled | n.s. | $F_{(1, 13)} = 12.36, p = 0.0038$ | n.s. |
| Acrophase | n.s. | $F_{(1, 13)} = 6.795, p = 0.0217$ | n.s. |
| Intradaily variability | n.s. | $F_{(1, 13)} = 5.613, p = 0.0340$ | n.s. |
| Free-running period | n.s. | n.s. | n.s. |
| 6-h phase advance | n.s. | n.s. | n.s. |
| 6-h phase delay | n.s. | n.s. | n.s. |
| Motor test |  | Two-way ANOVA |  |
|  | Sex | Viral vector | Interaction |
| Horizontal bar test | n.s. | $F_{(1, 29)} = 70.96, p < 0.0001$ | n.s. |
| Pole test | n.s. | $F_{(1, 29)} = 68.15, p < 0.0001$ | n.s. |
| Rotarod test | n.s. | $F_{(1, 29)} = 22.78, p < 0.0001$ | $F_{(1, 29)} = 5.889, p = 0.0217$ |

Concomitant to low activity levels in KO mice was a reduced rhythm stability and a shift of the central phase of the 24-h activity cycle. Particularly, intradaily variability was increased and the acrophase was delayed in KO mice (Fig. 1D, upper mid and right panel, Table. 1), which was also demonstrated by activity patterns (Fig. 1B).

Affective behaviours assessed in the open field and sucrose preference test were unchanged in KO animals compared controls (Suppl. Fig. 1A-C), suggesting that the knockout of *Bmal1* from the LHb did not cause behavioural abnormalities in general. However, an effect of the viral vector on the immobility time in the tail suspension test was found (F(1, 29) = 5.029, P = 0.0327, two-way ANOVA) indicating that KO mice move less when suspended by their tails (Suppl. Fig. 1D).

### Deletion of *Bmal1* from the LHb attenuates motor function and blunts oscillations of striatal DA levels and clock gene expression

Due to low wheel running activity observed in mice with habenular *Bmal1* ablation, motor functioning was furthermore evaluated using the horizontal bar test, pole test and rotarod test. All tests demonstrated a significant deficit in motor function in KO mice. Specifically, male and female KO mice performed worse in the horizontal bar and pole test compared to controls (Fig. 2A, left and mid panel, Table. 1). While the type of viral vector was decisive for rotarod performance, only KO females executed the test significantly worse compared to controls (Fig. 2A, right panel, Table. 1). Because motor functioning is modulated through the midbrain DAergic system, DA levels in tissue collected from the dorsal striatum at various times of the day were assessed using HPLC in a follow-up experiment.

**Figure 2.**
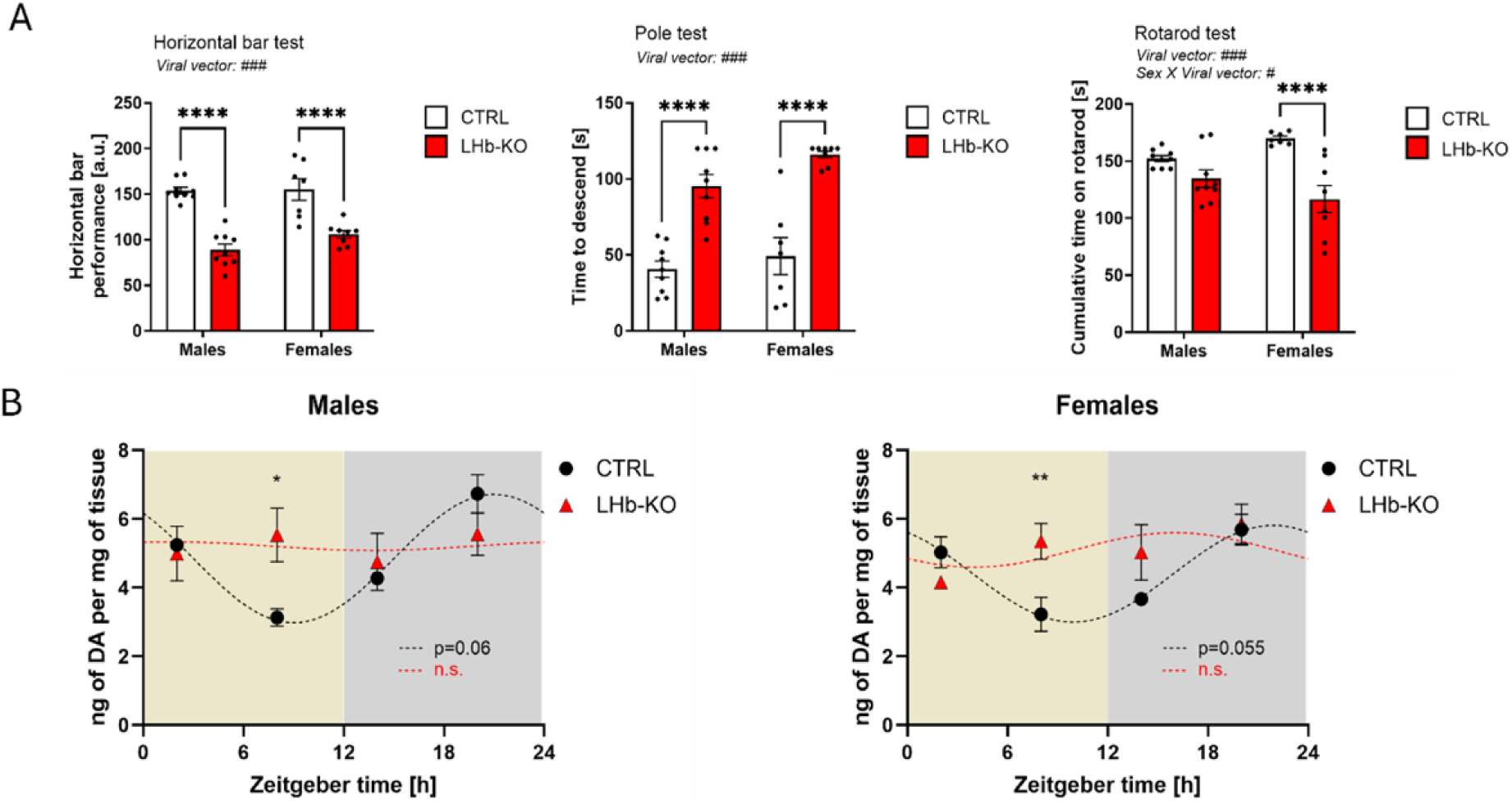
Striatal DA levels and motor functioning in LHb Bmal1 KO mice. A) Two-way ANOVAs were run for the: horizontal bar test, pole test, and rotarod. Males: n = 18 (9 KO); females: n = 15 (8 KO). B) Two-way ANOVAs were run for liquid chromatography coupled mass spectrometry measurements of striatal dopamine levels. Males and females: n = 6 (3 KO). Note: *p< .05; ** p<.01; ****p<.0001

Control mice displayed robust diurnal changes in DA levels peaking around the end of the dark phase (Fig. 2B). One-way ANOVA revealed a significant effect of daytime on DA levels in both CTRL males and females (Table. 2). Subsequent cosine analysis confirmed diurnal fluctuations of DA levels, although the results of zero-amplitude testing approached levels of statistical significance only (Table. 3). In contrast, KO animals displayed considerably blunted oscillations of DA levels across the day (Fig. 2B), which was confirmed by statistical analysis (Table. 3).

**Table 2.** Results of statistical analysis of daily striatal dopamine fluctuations in CTRL and KO animals.

| Sex | Viral vector | One-way ANOVA |  |
| --- | --- | --- | --- |
|  |  | Time point |  |
| Males | CTRL | $F_{(3, 8)} = 23, p = 0.0003$ | |
|  | KO | n.s. |  |
| Females | CTRL | $F_{(3, 8)} = 7.86, p = 0.009$ | |
|  | KO | n.s. |  |
| Sex | Viral vector | Two-way ANOVA |  |
|  |  | Time point | Interaction |
| Males | n.s. | $F_{(3, 16)} = 3.914, p = 0.0285$ | $F_{(3, 16)} = 3.395, p = 0.0438$ |
| Females | n.s. | $F_{(3, 16)} = 3.910, p = 0.0286$ | $F_{(3, 16)} = 3.527, p = 0.0392$ |

**Table 3.** Cosine analysis of diurnal changes in striatal dopamine levels measured by HPLC.

| Sex | Viral vector | Mesor | Amplitude | Acrophase (ZT) | Zero Amplitude test |  |  |
| --- | --- | --- | --- | --- | --- | --- | --- |
|  |  |  |  |  | F | P | R<sup>2</sup> |
| Males | CTRL | 4.85 | 1.87 | 20.99 | 115.8 | 0.065 | 0.995 |
|  | KO | 5.2 | 0.12 | 1.74 | 0.03 | 0.96 | 0.06 |
| Females | CTRL | 4.4 | 1.41 | 21.92 | 161.4 | 0.055 | 0.997 |
|  | KO | 5.1 | 0.5 | 16 | 0.246 | 0.818 | 0.33 |

Two-way ANOVA (factors: time of day, viral vector) furthermore verified that in both males and females, time of day was significantly affecting levels of DA in the dorsal striatum (Table. 2). Importantly, a significant interaction between time of day and the type of viral vector was found (Table. 2), further indicating that diurnal fluctuations in striatal DA depend on the presence of *Bmal1* in the LHb.

To decipher the consequences of disrupted striatal DA oscillations on MSN function, gene expression analysis was performed. Because previous work has established a link between DA signalling and circadian clock gene expression, daily profiles Bmal1, Per1 and Per2 were examined. Control males and females displayed robust diurnal fluctuations in striatal clock gene expression, whereas expression profiles were blunted in animals with a deletion of Bmal1 from cells of the LHb (Fig. 3). Statistical analysis by two-way ANOVA revealed a significant effect of daytime on expression profiles, although a significant interaction between time of day and viral vector indicated that oscillations of striatal clock gene expression were dependent on habenular Bmal1 expression (Table. 4).

**Figure 3.**
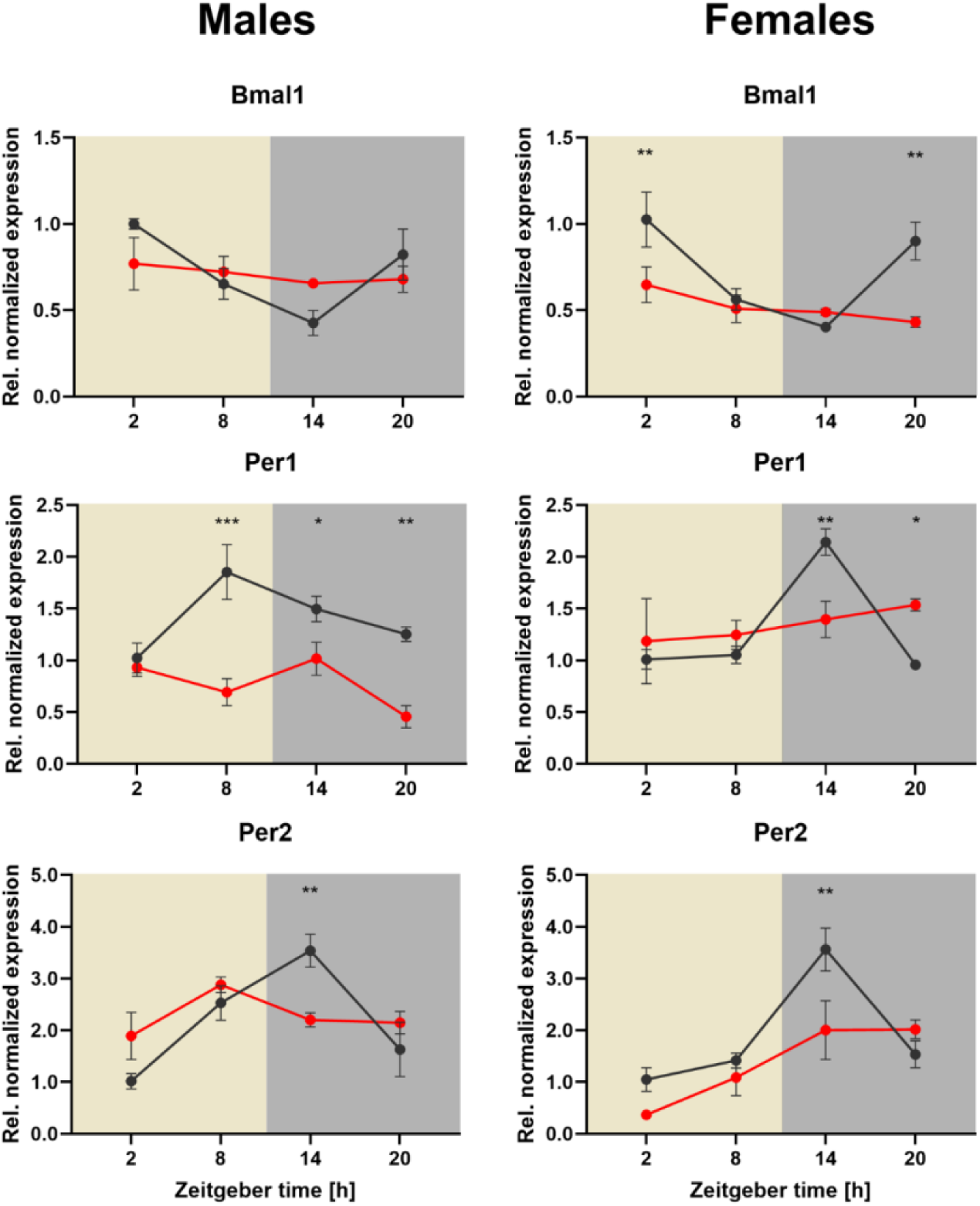
DS clock gene expression is blunted in KO mice. Two-way ANOVAs were run for all gene expression data, viral vector x time. Males: n = 6 (3 KO); females: n = 6 (3 KO). Top row: daily Bmal1 levels in males (left) and females (right). Middle row: Per1 rhythms in males (left) and females (right). Bottom row: Per2 rhythms males (left) and females (right). Note: *p< .05; ** p<.01; ***p<.001

**Table 4.** Results of statistical analysis of striatal gene expression profiles.

| Gene | Sex | Two-way ANOVA |  |  |
| --- | --- | --- | --- | --- |
|  |  | Viral vector | Time point | Interaction |
| *Bmal1* | Males | n.s. | $F_{(3, 16)} = 4.498, p = 0.018$ | n.s. |
| | Females | $F_{(1, 16)} = 11.18, p = 0.0041$ | $F_{(3, 16)} = 7.776, p = 0.002$ | $F_{(3, 16)} = 4.646, p = 0.0161$ |
| *Per2* | Males | n.s. | $F_{(3, 16)} = 9.035, p = 0.001$ | $F_{(3, 16)} = 4.841, p = 0.0139$ |
| | Females | $F_{(1, 16)} = 5.452, p = 0.0329$ | $F_{(3, 16)} = 15.73, p \leq 0.0005$ | $F_{(3, 16)} = 3.593, p = 0.0371$ |
| *Per1* | Males | $F_{(1, 16)} = 37.24, p \leq 0.0005$ | $F_{(3, 16)} = 4.016, p = 0.0262$ | $F_{(3, 16)} = 4.844, p = 0.0139$ |
| | Females | n.s. | $F_{(3, 16)} = 5.927, p = 0.0064$ | $F_{(3, 16)} = 4.936, p = 0.0130$ |

### Blunted striatal DA levels in mice with *Bmal1* deletion for the LHb are associated with changes in genes expression in the substantia nigra

Because extracellular DA levels in the dorsal striatum are regulated by DArgic neurons of the substantia nigra pars compacta, gene expression levels of circadian clock genes and tyrosine hydroxylase were assessed. While marked diurnal changes in of *Bmal1*, *Reverb-a* and *Th* expression were observed in both male and female control mice, levels of gene transcripts were blunted in KO mice (Fig. 4). Statistical analysis furthermore demonstrated that the presences of *Bmal1* in the LHb is a critical factor in the regulation of daily changes in *Bmal1* and *Th* expression in the substantia nigra (Table. 5). Although visual analysis of *Reverb-a* expression profiles indicated an impact of habenular *Bmal1* deletion, only time of day had a significant effect on diurnal levels of *Reverb-a* transcripts (Table. 5).

**Figure 4.**
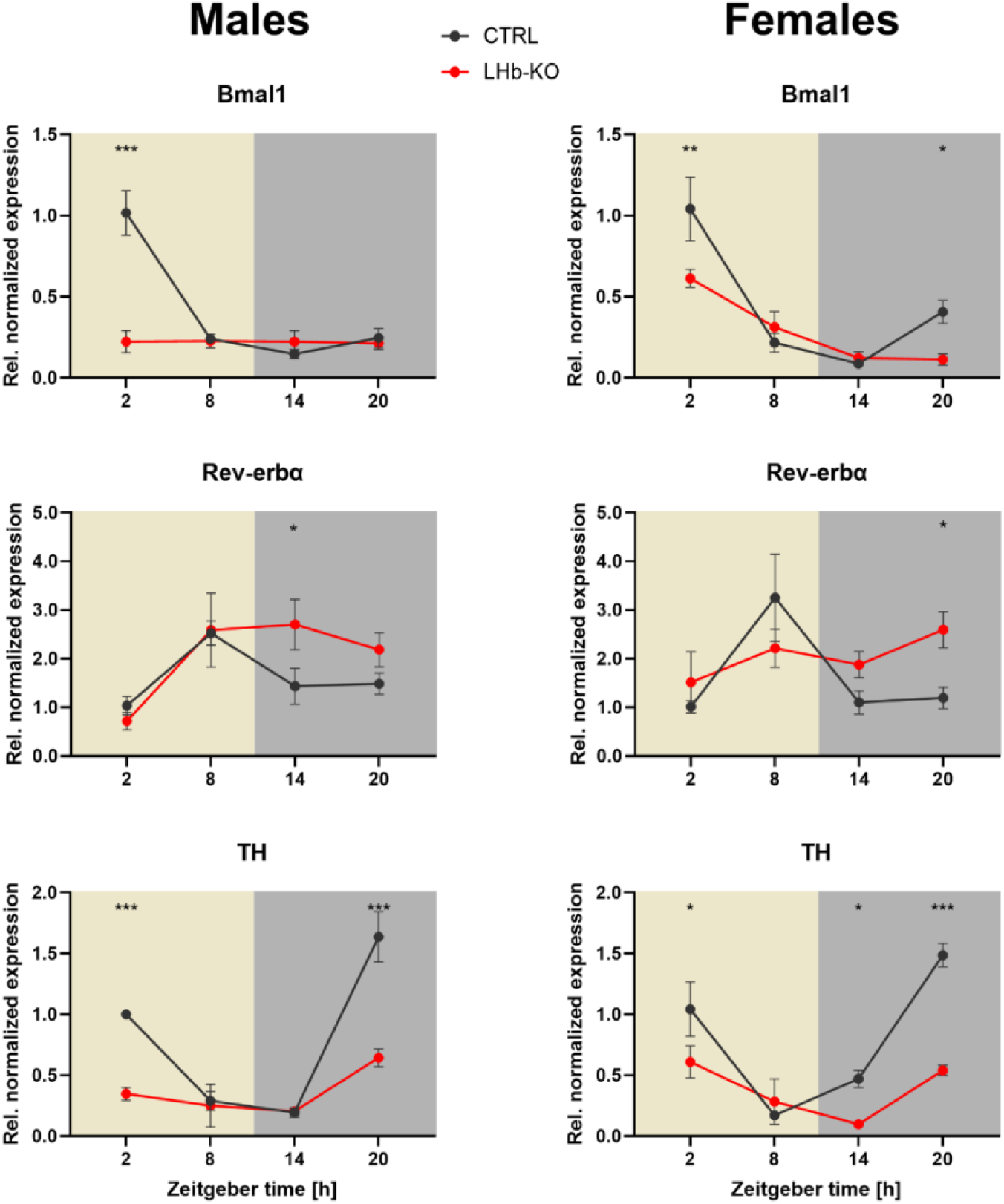
LHb Bmal1 KO altered SN gene expression. Two-way ANOVAs were run for all gene expression data, viral vector x time. Males: n = 6 (3 KO); females: n = 6 (3 KO). Top row: daily Bmal1 levels in males (left) and females (right). Middle row: Rev-erb⍺ rhythms in males (left) and females (right). Bottom row: Tyrosine hydroxylase rhythms in males (left) and females (right). Note: *p< .05; ** p<.01; ***p<.001

**Table 5.**
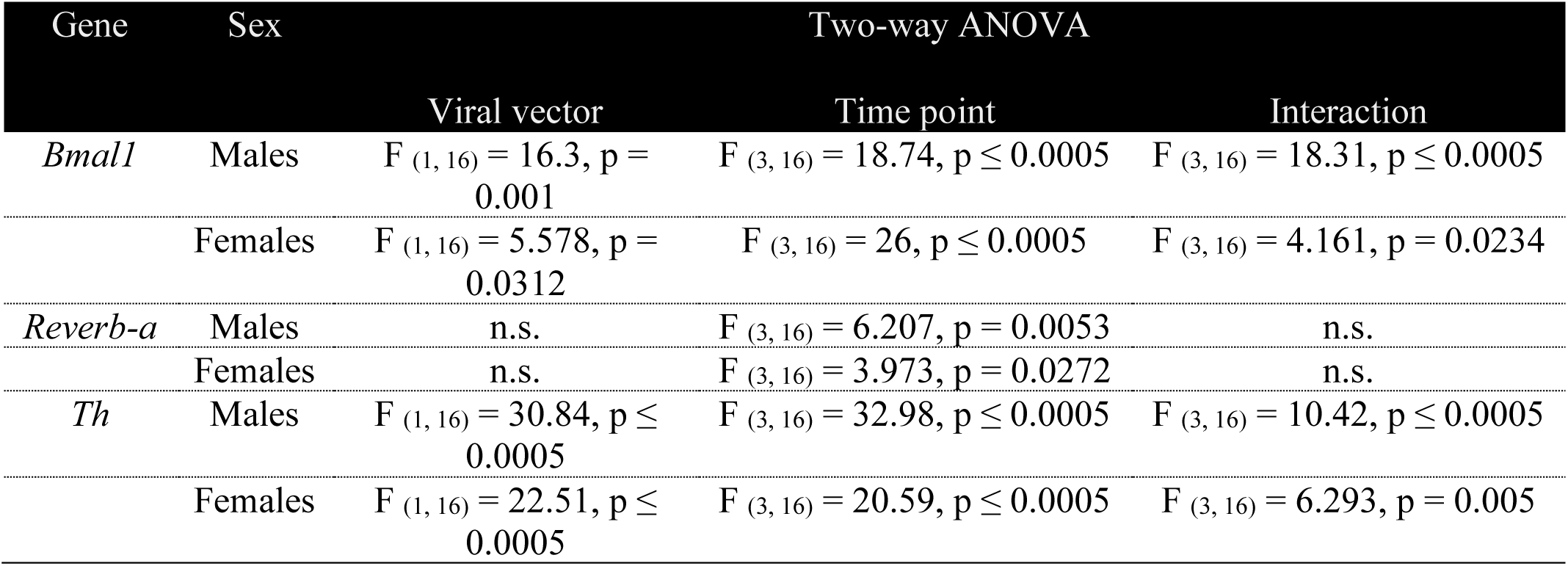
Results of statistical analysis of substantia nigra gene expression profiles.

| Gene | Sex | Two-way ANOVA |  |  |
| --- | --- | --- | --- | --- |
|  |  | Viral vector | Time point | Interaction |
| *Bmal1* | Males | $F_{(1, 16)} = 16.3, p = 0.001$ | $F_{(3, 16)} = 18.74, p \leq 0.0005$ | $F_{(3, 16)} = 18.31, p \leq 0.0005$ |
| | Females | $F_{(1, 16)} = 5.578, p = 0.0312$ | $F_{(3, 16)} = 26, p \leq 0.0005$ | $F_{(3, 16)} = 4.161, p = 0.0234$ |
| *Reverb-a* | Males | n.s. | $F_{(3, 16)} = 6.207, p = 0.0053$ | n.s. |
| | Females | n.s. | $F_{(3, 16)} = 3.973, p = 0.0272$ | n.s. |
| *Th* | Males | $F_{(1, 16)} = 30.84, p \leq 0.0005$ | $F_{(3, 16)} = 32.98, p \leq 0.0005$ | $F_{(3, 16)} = 10.42, p \leq 0.0005$ |
| | Females | $F_{(1, 16)} = 22.51, p \leq 0.0005$ | $F_{(3, 16)} = 20.59, p \leq 0.0005$ | $F_{(3, 16)} = 6.293, p = 0.005$ |

### Global rhythm disruption causes motor phenotypes resembling effects of *Bmal1* deletion from the LHb

Although various studies demonstrated a link between altered DA signalling and deficits in motor functioning in mice, effects of chrono disruption on motor performance are currently unknown. Blunted oscillations of striatal DA levels in KO mice were accompanied by attenuated performance in motor tasks suggesting functional causality, which is further supported by results in mice with abolished 24-h rhythms. Mice kept under prolonged time in constant light lost 24-h rhythmicity (Fig. 5C & D) and performed worse in the rotarod and pole test compared to animals kept under a standard light/dark cycle (Fig. 5 A&B). Specifically, time to descend was increased in chrono disrupted males and females compared to controls (Fig. 5A), while performance on the rotarod was reduced (Fig. 5B). Three-way ANOVA revealed a significant effect of the light condition and test phase on the performance in both tasks, and a significant interaction between these two factors indicated that indeed the prolonged exposure to constant light was causal to the observed motor deficits (Table. 6).

**Figure 5.**
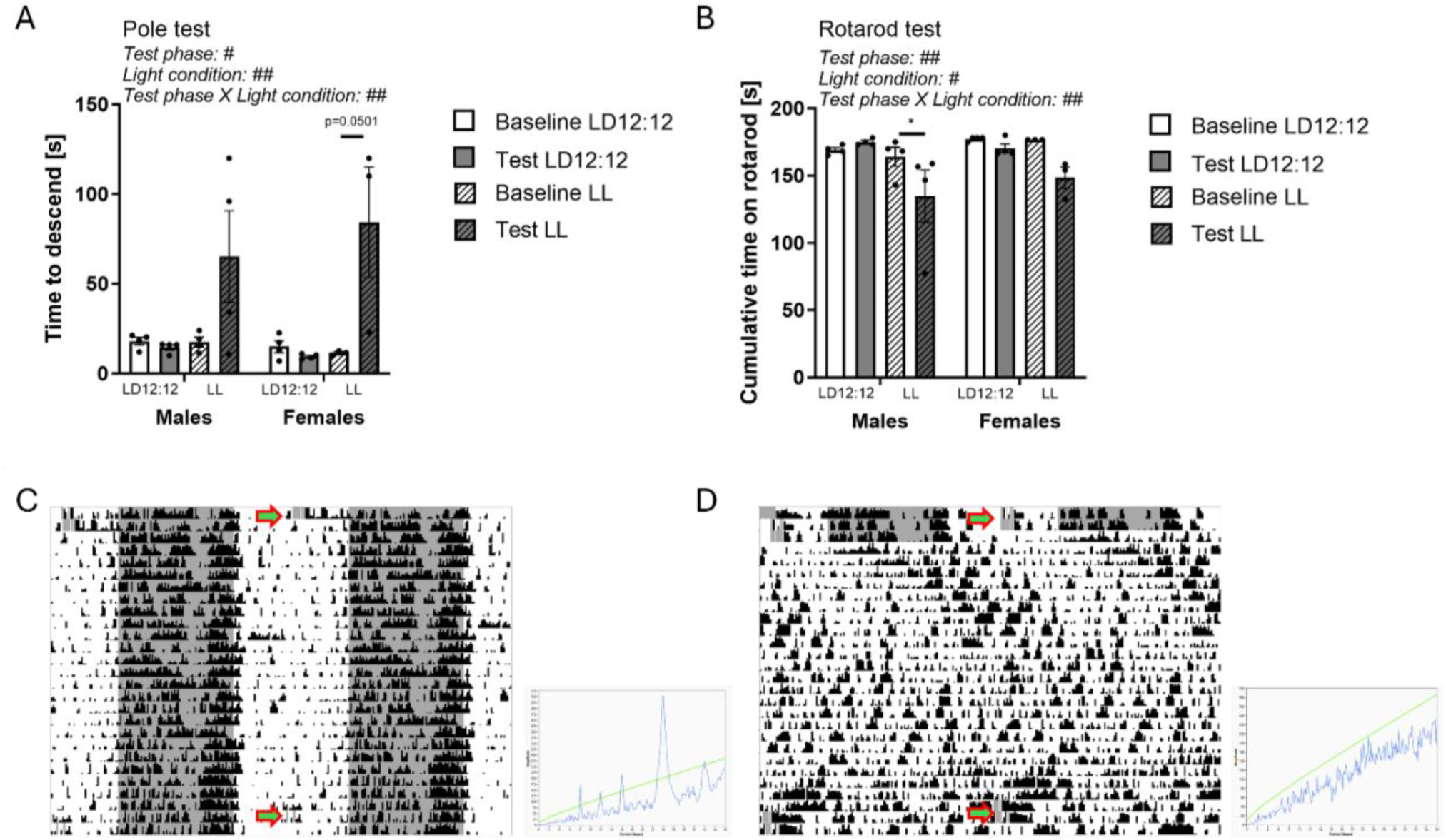
Mice under constant light performed worse on motor tests. Males: n = 8 (4 LL); females: n = 7 (3 LL). A) Two-way ANOVA of pole test performance before and after constant light exposure, light x time. B) Two-way ANOVA of rotarod performance at baseline and after constant light exposure, light x time. Males: *p< .05. C) Sample actogram of a wild-type mouse under typical 12:12 LD cycle. Arrows depict motor testing. D) Actogram of wild-type mouse kept under constant light for 3 weeks. Arrows show times of behavioural testing.

**Table 6.** Statistical analysis of motor performance in chronodisrupted animals.

| Motor test | Three-way ANOVA |  |  |  |
| --- | --- | --- | --- | --- |
|  | Sex | Test phase | Light condition | Interaction |
| Pole test | n.s. | $F_{(1, 11)} = 9.278$,<br>$p = 0.0111$ | $F_{(1, 11)} = 10.59$, $p = 0.0077$ | Test phase X Light condition:<br>$F_{(1, 11)} = 12.52$, $p = 0.0046$ |
| Rotarod test | n.s. | $F_{(1, 11)} = 14.48$,<br>$p = 0.0029$ | $F_{(1, 11)} = 5.18$, $p = 0.0439$ | Test phase X Light condition:<br>$F_{(1, 11)} = 13.27$, $p = 0.0039$ |

## DISCUSSION

Daily changes in brain and neuronal network functions are critical for the regulation of physiology and behaviour. Although various parts of the brain display diurnal rhythms, it has been shown that these oscillations dampen rapidly in the absence of rhythmic input (Begemann et al., 2020; Landgraf et al., 2016b). Few brain regions besides the SCN, such as the lateral habenula (LHb), have autonomous or semi-autonomous clocks enabling persistent oscillation independently of temporal cues (Guilding & Piggins, 2007a; Salaberry et al., 2019). Therefore, it has been proposed that the purpose of an extra-SCN oscillator in the LHb is to act as a circadian pacemaker for midbrain dopamine (DA) nuclei, thus driving daily oscillations in DAergic functions – which are pivotal for proper behavioural output, including mood, reward and motor functioning (Becker-Krail et al., 2022b; Pradel et al., 2022b; Proulx et al., 2014). The results of this study support this view. The conditional knockout of *Bmal1* in the LHb dysregulated daily oscillations of *Bmal1*, *Reverb-a,* and *Th* expression in the substantia nigra (SN), which was associated with an absence of daily DA oscillations and rhythms of clock gene expression in the dorsal striatum (DS). These outcomes suggest that the LHb is the pacemaker for the nigrostriatal pathway, synchronizing the striatal clock through daily changes in tonic DA release from the SN, thereby regulating motor functioning in mice.

The significance of the relationship between the circadian system and midbrain DAergic circuits has been recognized in various studies (Becker-Krail et al., 2022a; Pradel et al., 2022a). Rhythm analysis in isolated midbrain DAergic nuclei, however, demonstrated that most of these nuclei have weak circadian clocks characterized by rapidly dampening oscillations in the absence of rhythmic input (Landgraf et al., 2016a). Because the central pacemaker in the SCN does not innervate midbrain DAergic nuclei directly, it has been suggested that temporal cues must be relayed through extra-SCN oscillators in the brain (Guilding & Piggins, 2007b), including from the LHb. Despite some discrepancy in the literature regarding its intrinsic rhythmicity (Guilding & Piggins, 2007a; Landgraf et al., 2016a; Salaberry et al., 2019), the LHb has been classified as a semi-autonomous oscillator receiving indirect SCN and retinal input (Baño-otálora & Piggins, 2017; Young et al., 2022a). Notably, a link between LHb clock gene expression and its molecular and neurophysiological properties, especially in mood and reward-related processed has been established in the past (Li et al., 2021a; Mendoza, 2017; Olejniczak et al., 2020; Sakhi et al., 2014; Salaberry et al., 2019; Young et al., 2022a). However, most of these studies have been conducted in global clock gene KO animals, which limits the conclusion that can be made about a direct effect of the LHb clock on the regulation of downstream brain regions. As such, the present study elected to use a targeted LHb- specific *Bmal1* KO to circumvent this issue.

Because the deletion of *Bmal1* within the molecular clockwork confers to loss in rhythmicity of cellular functions including gene expression (Bunger et al., 2000), it can be concluded that the LHb clock in *Bmal1* KO mice is non-functional. Importantly, the results of running wheel locomotor activity in the present study demonstrate that the conditional *Bmal1* knockout did not affect central clock function. Central clock properties, like free-running period and phase responses following advancing and delaying shifts of the light/dark cycle were unaffected by the KO. These outcomes therefore suggest that irregularities of diurnal rhythms in LHb efferent regions directly or indirectly, must stem from the LHb *Bmal1* knockout itself.

Despite the role of circadian clock genes *Per1* and *Per2* in the LHb in the regulation of affective behaviours (Li et al., 2021b; Olejniczak et al., 2021; Young et al., 2022b), only a weak association was found regarding the contributions of *Bmal1* in the present study. Mice with a conditional *Bmal1* knockout displayed slightly higher depressive-like behaviour in the tail suspension test compared to controls, whereas anhedonia-like behaviour was unchanged in the sucrose preference test. The overall lack of an aberrant affective phenotype follows similar literature where a conditional knockout of *Bmal1* in the forebrain did not affect anxiety-like behaviour in mice (Price et al., 2016), nor did targeted deletion of Bmal1 in the striatum impact performance on these same tests (Schoettner et al., 2022). Unlike deletions of *Bmal1*, clocks with single knockouts of *Per1* or *Per2* retain rhythmicity (Zheng et al., 2001). These outcomes therefore suggest that individual gene effects of *Per1* and *Per2* within the LHb account for changes in mood-related behaviours, conceivable through their impact on neuronal excitability (Young et al., 2022b), while dysregulation of the habenular clock through *Bmal1* deletion has only minor impacts on mood and affect.

*Bmal1* KO mice displayed strong deficits in motor functioning. Wheel running activity was markedly reduced in males and females after a deletion of *Bmal1* in the LHb and this was then supported by poor performance in the rotarod, horizontal bar, and pole tests. However, it is important to note that spontaneous activity assessed in the open field test was unaltered in KO animals, suggesting normal baseline motor functioning. The authors of a study on the genetic ablation of the dorsomedial habenula, which revealed almost identical motor phenotypes, argued that impairments in both, wheel running activity and rotarod performance are due to a lack of motivation rather than a motor deficit (Hsu et al., 2014). The results of the present study, however, contradict this view. Sucrose preference in *Bmal1* LHb KO mice was unchanged despite a prolonged fasting period prior to the test, indicating similar levels of motivation in KOs and controls. Moreover, the rotarod, pole and horizontal bar tests are measures of motor coordination and execution, and their potential association with motivated behaviours is inconsistent (Campos et al., 2013; Liebetanz et al., 2007). As such, it can be concluded that factors other than simply a lack of motivation must therefore contribute to the motor deficits found in *Bmal1* KO mice.

Consistent with the results of the present study, a reduction of wheel running activity has been observed in rats treated with 6-hydroxydopamine (6-HODA) in the medial forebrain bundle (Hood et al., 2010). Similarly, there is clear evidence showing the direct effects of DA deficiency on rotarod and pole test (Glajch et al., 2012; Ma & Rong, 2022; Ogawa et al., 1985; Sedelis et al., 2001), indicating that altered levels of extracellular DA in the striatum may be causal to the observed motor phenotypes. Unlike KO animals, mice with unilateral 6-HODA lesions or MPTP treatment, which results in a loss of striatal dopamine, typically display reduced spontaneous activity in the open field test (Fredriksson et al., 2011; Slézia et al., 2023). However, DA levels measured in the dorsal striatum in KO mice were not reduced per se, but their amplitude was markedly decreased, indicating that daily oscillations in DA tone were blunted. Indeed, numerous studies have supported the presence of daily DA oscillations in midbrain DAergic circuits, particularly in the striatum, and it has been suggested that a change in DA tone across the day has implications in the regulation of DAergic functions (Pradel et al., 2022). The outcomes of the motor tests conducted in the present study therefore indicate that a dysregulation of daily DA oscillations, rather than a loss of DA, may be sufficient to attenuate motor functioning in *Bmal1* KO mice. Indirect support for this hypothesis was provided by the outcomes of motor tests conducted in arrhythmic animals, showing that a systemic dysregulation of circadian rhythms can impact motor functioning.

Unknown to date is the underlying mechanism leading to motor dysfunction in KO mice. Rhythms of clock gene expression and DA in the DS were blunted in KO mice. Because DA signaling affects clock gene expression in the striatum (Hood et al., 2010; Imbesi et al., 2009), it is tempting to speculate that daily changes in DA tone could be a zeitgeber to the striatal clock, which rapidly dampens in the absence of external cues (Landgraf et al., 2016a). This, in turn, would indicate that the loss of rhythmicity in striatal DA tone may be causal for blunted rhythms of clock gene expression in the striatum. Interestingly, results from previous studies suggest that circadian clock gene expression in the striatum contributes to the regulation of motor function in mice (Schoettner et al., 2022). Ablation of *Bmal1* from medium spiny neurons (MSNs) of the striatum was associated with strongly attenuated motor performance in the horizontal bar and rotarod test. This effect appeared to be limited to the deletion of *Bmal1*, as motor performance was unaffected in mice with a conditional knockout of *Per2* (Schoettner et al., 2022). However, while motor coordination was affected by the conditional *Bmal1* knockout, voluntary wheel running activity was unchanged and spontaneous activity assessed in the OFT was increased (de Zavalia et al., 2021; Schoettner et al., 2022). The discrepancy in these results could be attributed to the differences in the levels of striatal *Bmal1* expression. While mice with striatum-specific *Bmal1* KO have no functional copies of the gene in MSNs (Schoettner et al., 2022), *Bmal1* was still expressed in KO mice here. Although it has been shown that loss-of-function of *Bmal1* in the striatum affected behaviour differently compared to its downregulation (Porcu et al., 2020), future experiments have to clarify the significance of potential dose effects of *Bmal1* in MSN in the context of motor functioning.

Alternatively, alterations in motor functioning, such as wheel running activity in *Bmal1* KO mice, may stem from changes in signaling pathways other than midbrain DAergic systems. The LHb also innervates serotonergic (5-HT) nuclei such as the dorsal raphe nucleus (DRN), which in turn projects to nuclei in the basal ganglia including the SN and the striatum (Huang et al., 2019; Reed et al., 2013). While an association between 5-HT and motor functioning has been established in the past (Weber et al., 2009), follow-up studies must elaborate the link between the LHb, 5-HT, and motor functioning in more detail.

In summary, this work provides compelling evidence that the LHb is a circadian pacemaker within the nigrostriatal pathway. By coordinating rhythms of DA synthesis in the SN, it regulates daily oscillations of striatal DA tone responsible for synchronizing the striatal circadian clock, which is critical to proper motor functioning in mice. While other monoaminergic systems must be considered in the regulation of striatal functioning and motor control, this study provides a basis for future investigations in the underlying molecular and neurophysiological mechanism and brain circuits regulating motor function.

## METHODS

### STAR METHODS

#### Key resource table

| REAGENT or RESOURCE | SOURCE | IDENTIFIER |
| --- | --- | --- |
| Antibodies |  |  |
| Rabbit polyclonal anti-Bmal1 (1:500) | Novus Biologicals | Cat# NB100-2288<br>RRID:AB_10000794 |
| Mouse monoclonal anti-NeuN (1:2000) | Abcam | Cat# ab104224<br>RRID:AB_10711040 |
| Donkey anti-Mouse IgG Alexa Flour 594 (1:500) | Thermo Scientific™ | Cat# A-21203<br>RRID:AB_141633 |
| Donkey anti-Rabbit IgG Alexa Flour 647 (1:500) | Thermo Scientific™ | Cat# A-31573<br>RRID:AB_2536183 |
| Bacterial and virus strains |  |  |
| AAV2/9-CAG-Cre-eGFP | Canadian Neurophotonics Platform Viral Vector Core Facility | Cat# 125-aav9<br>RRID:SCR_016477 |
| AAV2/9-CAG-eGFP | Canadian Neurophotonics Platform Viral Vector Core Facility | Cat# 475-aav9<br>RRID:SCR_016477 |
| Chemicals, peptides, and recombinant proteins |  |  |
| 2-(3,4-Dihydroxyphenyl)ethyl-1,1,2,2-d<sub>4</sub>-amine HCl | C/D/N ISOTOPES INC | Cat# D-1540<br>CAS# 203633-19-6 |
| Dopamine Hydrochloride | Thermo Scientific™ | Cat# AAA1113606<br>CAS# 62-31-7 |
| Methanol HPLC Grade | Millipore Sigma | Cat# 34885<br>CAS# 67-56-1 |
| Formic Acid | Sigma Aldrich | CAS# 64-18-6 |
| Acetonitrile | Sigma Aldrich | CAS# 75-05-8 |
| Ultrapure Water | Thermo Scientific™ | Cat# 10977015 |
| Cytiva HyClone™ Phosphate Buffered Saline | Fisher Scientific | Cat# SH3025601 |
| TRIzol™ Reagent | Thermo Scientific™ | Cat# 15596018 |
| DEPC-Treated Water | Thermo Scientific™ | Cat# AM9906 |
| Nuclease-Free Water | Thermo Scientific™ | Cat# AM9937 |
| Sodium Chloride | Bioshop | Cat# SOD002.10 |
| Sodium phosphate monobasic | BioShop Canada | Cat# SPM306 |
| Sodium phosphate dibasic | BioShop Canada | Cat# SPD307 |
| Polyvinylpyrrolidone | Sigma Aldrich | Cat# PVP-1 |
| Ethylene glycol | Sigma Aldrich | Cat# E9129 |
| Paraformaldehyde | Electron microscopy sciences | Cat#19202 |
| Triton™ X-100 | Sigma Aldrich | Cat# T8787 |
| Normal Donkey Serum | Sigma Aldrich | Cat# 566460 |
| Skim Milk Powder |  |  |
| ProLong™ Diamond Antifade Mountant | Thermo Scientific™ | Cat# P36971 |
| Ketamine Hydrochloride | Zoetis | CAS# 1867-66-9<br>DIN 02173239 |
| Xylazine Injection | Rompun | Lot# 20015100 |
| Anafen Injection | Merial | DIN 01938126 |
| Isoflurane | PPC | Lot# I0858II5 |
| Isopentane (2-Methylbutane) | Fisher Scientific | Cat# O3551-4<br>CAS# 78-78-4 |
| Critical commercial assays |  |  |
| iScript™ Reverse Transcription Supermix | BioRad | Cat# 1708841 |
| SsoAdvanced Universal SYBR® Green Supermix | BioRad | Cat# 1725274 |
| Pierce BCA Protein Assay Kit | Thermo Scientific™ | Cat# 23227 |
| Experimental models: Organisms/strains |  |  |
| B6.129S4(Cg)-Arntl<sup>tm1Weit</sup>/J | The Jackson Laboratory | RRID: MGI:6275181 |
| Oligonucleotides |  |  |
| Primers for quantitative real-time PCR, see Table SX | This paper | N/A |
| Software and algorithms |  |  |
| Actitrack 2.7.13 | Panlab | https://www.panlab.com/ |
| Bio-Rad CFX Maestro 2.0 | Bio-Rad | https://www.biorad.com/ |
| Clocklab 6 | Actimetrics | https://www.graphpad.com/ |
| ImageJ | ImageJ software | https://imagej.net/ |
| Prism 10 | GraphPad Software | https://www.graphpad.com/ |
| Stopwatch+ | Center for Behavioral Neuroscience, Atlanta, GA, USA | https://cbn-atl.org/ |
| Xcalibur™ | Thermo Scientific™ | https://www.thermofisher.com/ |
| Other |  |  |

#### Subjects

*Bmal1* floxed mice (B6.129S4(Cg)-Arntl^tm1Weit^/J) were originally obtained from Jackson Laboratories. Male and Female mice were group-housed under a 12:12 hrs light-dark (LD) cycle at 21 ± 2°C and 60% relative humidity with food and water available *ad libitum* prior to the stereotactic delivery of viral vectors at 12 – 16 weeks of age. The bedding of the cages was changed every week. Animals were kept individually after the surgery under the same housing conditions as described above unless stated otherwise.

This study was conducted following the guidelines and requirements of the Canadian Council on Animal Care (CCAC), approved by the Concordia University ethics committee (AREC number 3000256).

#### Stereotactic Surgery

Mice were anesthetized by intraperitoneal injection of Ketamine (100 mg/kg body weight) and Xylazine (10 mg/kg body weight) solution and received subcutaneous administration of Ketoprofen (5 mg/kg body weight) as post-operational analgesia. Animals were placed in a stereotaxic apparatus (KOPF, Tujunga, CA, USA) and received bilateral microinjections of recombinant viruses in the LHb (AP: -1.65, ML: 0.4, and DV: -3.1) using a 30-gauge needle attached to a 10 ul Hamilton syringe which was connected to a micropump (Pump 11 Pico Plus, Harvard Apparatus, Holliston, MA, USA). The injector was inserted at a 0° angle and the virus was delivered at a rate of 100 nl/min (150 nl total volume). For the generation of conditional knockout animals, recombinant viral vectors expressing Cre and eGFP (AAV2/9-CAG-Cre-eGFP, 1x10^12^ vg/ml) were injected. Control animals received viruses expressing eGFP only (AAV2/9-CAG-eGFP, 1x10^12^ vg/ml). The injector was left in place for five minutes following the injection to optimize diffusion. Mice were allowed to recover for three weeks before behavioural testing. LHb-specific expression of viral vectors was evaluated at the end of the experiment. Animals with missing, incomplete or off-region eGFP expression were excluded from all experiments.

#### Open Field Test

The open field test was used to assess anxiety-like behaviour and motor activity. The test was conducted at Zeitgeber time 2 (ZT2, ZT0 represents the time of lights-on). After a 30-minute habituation period in the experimental room, mice were placed in the corner of the open field arena (45 x 45 x 60 cm, Panlab, Barcelona, Spain) facing the wall. The open field was equipped with infrared beams to track horizontal and vertical activity over 60 minutes using the ACTItrack software (Panlab, Barcelona, Spain). At the end of the session, animals were weighed and returned to home cages while the arena was wiped with 70% ethanol solution (v/v in tap water) to remove any residues and olfactory cues before the next set of animals were tested. Total distance travelled, speed, resting time, rearing behaviour (vertical movements) as well as permanence time in the central and peripheral area of the open field and the latency to enter the central area was assessed.

#### Sucrose Preference Test

The sucrose preference test for assessing anhedonia in laboratory animals as a proxy of depressive-like behaviour. One week prior to the beginning of the experiment, animals were habituated to two drinking bottles containing tap water (50 ml Falcon tubes with sipper). To minimize the effects of neophobia as a confounding factor, animals were given two bottles of 1% sucrose solution (w/v in tap water) for 3 hours the day before testing, and the volume of consumed sucrose solution was assessed to ensure all animals were drinking. Food and water were removed from animal cages 14 hours before the sucrose test was conducted to increase the motivation to consume sucrose solution. Two hours after the beginning of the light phase on the test day (ZT2), food was returned and one bottle of water and one bottle of 1% sucrose solution were inserted into the cage. The bottles were left in the cage for one hour, at which time they were removed and weighed to determine the amount of sucrose consumed (g/kg body weight) and the preference for sucrose solution (Vsucrose solution/(Vsucrose solution + Vwater)).

#### Tail Suspension Test

In the tail suspension test, escape-related behaviour is quantified as a proxy for depressive-like phenotypes in mice. The apparatus consists of a metal stand on which mice were suspended by their tails while being videorecorded (Samsung Galaxy A5 2017) for 6 minutes. Animals were habituated to the conditions of the experimental room for 30 min before the beginning of the test, which was conducted at ZT8. To prevent mice from tail climbing during the test, climb stoppers were placed over the proximal part of the tail (Can et al., 2011). After completion of the 6 minutes trial period, animals were weighed and returned to their home cages. Two experienced researchers blind to the experimental conditions analyzed the video recordings using an on-screen stopwatch (Stopwatch+, Center for Behavioural Neuroscience, Atlanta, GA, USA). The total amount of mobility time (in seconds), defined as any movement of the body, was scored and subtracted from 360 seconds to determine immobility time.

#### Horizontal Bar Test

Motor coordination was assessed in a horizontal bar test. The experimental setup comprised a squared cardboard box (41 cm × 41 cm × 41 cm) with a metal bar (ø 2.5 or 5 mm) mounted on the top center of two opposite sides of the box. A cotton soft pad was placed on the bottom of the box to cushion animals falling from the bar. The test was conducted between ZT6 – 8 following a 30-minutes period to habituate animals to the experimental room. During the first stage of the test, mice were gently raised by their tail to the center of the 2.5 mm bar, which they had to grasp with the forepaws before being released. Within a 60 seconds trial period, mice had to either transverse the rod and touch one side of the box with a paw, stay on the rod for 60 seconds, or fall, while being video recorded (Samsung Galaxy A5 2017). The test was repeated three times with a 30 second interval between each trial, before it was conducted on the thicker rod (ø 5 mm) following the same experimental design. Videos were analyzed using a stopwatch and a score was calculated for each trial based on the mice’s performance, i.e. the time to reach one of the criteria mentioned above. For transversing the bar, the score was calculated by subtracting the time reaching the end of the bar from 120 (score pass = 120 - time pass). The score for falling represented the same time it took the mice to fall from the bar (score fall = time fall). If a mouse stayed on the bar for the entire 60 second trial period, a score of 60 was given. A cumulative score from the averages of the three trials of each experimental stage was calculated.

#### Pole Test

The pole test is common test to assess motor function in mice. Animals were habituated to the experimental room for 30 minutes prior to the test, which has been conducted between ZT8- ZT10. Mice were placed head-upward on the top of a vertical rough-surfaced pole (diameter 8 mm; height 55 cm) placed in an enclosure filled with bedding (5 cm) to cushion animals falling from the pole. Mice were trained on how to descend the pole before testing began. Mice were given 120 seconds to descend the pole, time was stopped upon reaching the enclosure surface. Scores were averaged over three trails, with 30 seconds intertrial intervals. If mice failed to turn downward and descend the pole or if the mouse fell, time was taken as 120 second (default value). The criteria assessed includes the proportion of those who were able to complete the turn and the total time to descend to the ground over three trials. Animals were video recorded (Samsung Galaxy A5 2017) and the videos were scored by an experienced researcher blind to the experimental conditions using a stopwatch.

#### Rotarod Test

Motor coordination was furthermore assessed on a custom-made, fixed-speed rotarod (Concordia University, Montreal, Canada). The rotarod was composed of a rod (ø 5 cm) mounted 19.5 cm above the base of the device, which was separated into 6 individual compartments equipped with soft foam pads to cushion mice falling off the rod. Two flanges (ø 20 cm) separated a 9.3 cm interspace on the rod, which was driven by an electric motor. A gearbox was used to set the desired turning speed of the rod. On the test day, animals were habituated to the experimental room 30 minutes before they underwent a training session on the rotarod between ZT2–4 to get familiar with the apparatus. Animals were gently lifted by their tail and carefully placed on the rod turning at 2 rpm. Animals falling from the rod were put back on the rod immediately throughout the 5-minute training session. Animals returned to their home cages until the actual test was conducted between ZT6-8 at the same day. Mice were placed on the turning rod as described above and left on the rod for a 1-minute trial while being videorecorded. In contrast to the training session, mice were not placed back on the rod within the trial if they fell off the rod unless it was due to poor placement by the experimenter. In that case the trial was not counted and repeated after a short resting time. The trial was repeated two more times with 30 second intertrial periods before the rod was set to a higher speed. The starting speed of 2 rounds per minute (rpm) was gradually increased to 4, 8 and 12 rpm. Animals that failed to stay on the rod for 1 minute in all three trials were not tested further at higher speeds. Rotarod performance was given as a cumulative score calculated from the average times of the 3 trials at each speed.

#### Locomotor Activity Rhythms

To verify that the circadian system and gross locomotor activity was not affected by the deletion of *Bmal1* from the LHb, 15 weeks old mice were housed individually in cages equipped with running wheels (Actimetrics, Wilmette, IL, USA) food and water were provided *ad libitum*.

Cages were placed in light-proof cabinets equipped with programmable lights (Actimetrics, Wilmette, IL, USA). At the beginning of the experiment, mice were kept under a standard 12:12 h light-dark (LD) cycle with the light intensity set to 200 lux during the light phase and 0 lux during the dark phase to assess basic measures of daily rhythms in locomotor activity.

Thereafter, mice were exposed to 6 h phase advance (+6h), followed by a 6 h phase delay (-6h) to evaluate the capacity of the circadian clock to adjust to phase changes of the LD cycle.

Intrinsic properties of the circadian clock were assessed in animals exposed to constant darkness (DD). Mice were kept for 21-40 days under each respective light regimen. Wheel running was recorded and analyzed using Clocklab 6 (Actimetrics, Wilmette, IL, USA). Basic measures such as mesor, amplitude, activity onset and offset, and rhythm stability was analyzed from summed activity counts (10-minute bins) recorded over a period of 10 days. The time to re-entrain to 6-h phase shifts was determined for activity onset and offset individually by calculating the number of days from the shift of the light cycle until animals were stably alignment to the shifted light cycle. The free-running period was determined using Chi-Square periodogram in Clocklab 6 and double-plotted actograms were prepared to visualize representative locomotor activity patterns. Effects of central clock disruption on motor functioning was studied in a subset of male and female mice exposed to constant light. Animals were kept individually under standard housing conditions to assess motor function (rotarod and pole test) before they were randomly assigned into two groups. While one group of animals continued to be kept under standard conditions, the second group was exposed to constant light (∼ 500 lx) for 3 weeks. Locomotor activity rhythms were assessed using infrared motion detectors mounted on top of the cages (Actimetrics, Wilmette, IL, USA) to determine the degree of chronodisruption of each animal visually and using the Chi^2^-peiodogram in Clocklab 6. For this, the last 10 days before motor testing were analyzed. Animals of all groups were re-tested on the pole and rotarod test once mice kept under LL had no detectable rhythms anymore.

#### Tissue collection

Fresh frozen tissue was used for gene expression and LC-MS analysis. For this, brains from animals live-decapitated at corresponding times of the day were rapidly extracted and submerged in Isopentane cooled down to -30 °C for 3 minutes and stored at -80 °C thereafter. Serial coronal sections (100 µm thick) were obtained using a Cryostat (Microm HM 505 E, Microm International, Walldorf, Germany). The dorsal striatum (DS), the substantia nigra (SN) and the LHb were identified using the mouse brain atlas (Paxinos & Franklin, 2003), and tissue punches of the DS (1.5 mm in diameter) and SN (1 mm in diameter) were collected from each hemisphere. Region-specific expression of recombinant viral vectors was evaluated in brain sections of the LHb. The samples were kept in -80 °C until further processing. Formaldehyde-fixed tissue was prepared for immunostainings. Animals were deeply anesthetized by exposure to an atmosphere of isoflurane and then transcardially perfused by cold saline (0.9% sodium chloride, pH 7.2) followed by paraformaldehyde solution (PFA, 4% in 0.1M phosphate buffer, pH 7.2) using an infusion pump. Brains were dissected and postfixed in PFA solution for 22 – 24h at 4°C thereafter. Coronal sections of brain tissue (30 um) were collected using a Leica vibratome (Leica Biosystems Inc., Concord, Ontario, Canada) and analyzed under a fluorescent microscope (Leica DM4000B, Leica Microsystems, Concord, Ontario, Canada) to validate region-specific viral vector delivery. Brain slices used for immunofluorescence imaging were stored at -20°C in Watson’s cryoprotectant (Watson et al., 1986).

#### Liquid Chromatography Mass Spectrometry

2-(3,4-Dihydroxyphenyl)ethyl-1,1,2,2-d4-amine HCl (D4-DA, C/D/N ISOTOPES) was used as an internal standard in the extraction solution. The DA (DA) standards were prepared using pooled caudate putamen tissues (four wildtype mice, half female, half collected at ZT8 half at ZT20) to final concentrations of 0.05, 0.1, 0.5 and 1 µM of labelled DA standard. The calibration curve was plotted using the ratios of peak areas of DA (DA isotope) to D4-DA (labelled isotope) and the slope was used to calculate DA concentrations in samples. Briefly, tissue extraction was done using frozen brain tissue samples were thawed on ice and homogenized in 150 ul of an 80% methanol and isotope-labelled DA (to a final concentration of 2uM) in ultrapure water (v/v). The mixture was centrifuged at 4 °C, 14,000 rpm, for 20 min. The supernatant was transferred into a new tube and dried under nitrogen stream on ice. The residue was reconstituted in 1ml of 0.1M 3NT standard in ultrapure water (v/v).

#### Liquid chromatography mass spectrometry (LC-MS) analysis

LC-MS analyses were performed on an Agilent 1100 LC system coupled to a Thermo LTQ Orbitrap Velos mass spectrometer (Thermo Scientific™, Waltham, MA, USA) equipped with a heated electrospray ion source at positive mode. A Waters Atlantis dC18 column (100 x 2.1 mm, 3 μ particle diameter, Waters) was used and target compounds were eluted using an 18-min gradient at a flow rate of 250 µL/min with mobile phase A (99.9 % ultrapure water and 0.1 % formic acid (FA)) and B (99.9 % Acetonitrile and 0.1 % FA). The gradient started at 2 % B and held for 4 min, linear gradients were achieved to 90 % B at 6 min, then followed by isocratic with 90 % B for 2 min and with 2 % B for 9 min. 5 µL of each sample was injected for the LC-MS analysis. MS spectra (*m/z* 100-250) were acquired in the Orbitrap at a resolution of 60000, DA and D4-DA were targeted at *m/z* 154.0863 and 158.1114 at retention time of 2.5 min. Peak area values were extracted using Thermo XCalibur software (v2.2 SP1.48).

#### Gene expression analysis

Total RNA was isolated from brain tissue of the DS and SN using a standard Trizol protocol according to the manufacturer’s instructions (Ambion, Carlsbad, CA) and RNA yield was measured using spectrophotometry (Nanodrop 2000; Thermo Scientific™, Wilmington, DE, USA). Integrity of the isolated RNA was assessed by the “bleach gel” method (Aranda et al., 2012) and cDNA was synthesized from 1 ug of RNA using the iScript^TM^ Reverse Transcription Supermix (Biorad, Hercules, CA, USA) thereafter. A no reverse transcriptase (no-RT) control was prepared along with the cDNA samples. Levels of gene expression were measured using SYBR green based quantitative real-time PCR. For this, 10 ul reactions containing SsoAdvanced Universal SYBR® Green Supermix (Biorad, Hercules, CA, USA) and 300 uM of the respective forward and reverse primers were amplified in triplicate using a CFX96^TM^ Real-Time PCR system (Biorad, Hercules, CA, USA). The list of primers is provided in Table 1. The CFX Maestro Software (Biorad, Hercules, CA, USA) and Microsoft Excel were used to calculate relative normalized changes in gene expression by the delta-delta Ct (ΔΔCt) method (Pfaffl, 2001). The levels of target gene expression relative to two reference genes were further normalized to the sample collected from CTRL animals at ZT2.

#### Immunofluorescence staining

Free-floating sections kept in Watson’s cryoprotectant were rinsed for 10 minutes in phosphate buffered saline (PBS, pH 7.4) followed by three rinses of 10 minutes each in a solution of PBS containing 0.3% Triton-X (PBST). The brain sections were kept in a blocking solution (3% skim milk powder, 6% normal donkey serum in PBST) for one hour and incubated in a PBST solution containing 3% skim milk powder, 2% normal donkey serum, rabbit Anti-BMAL1 primary antibody (1:500, NB100-2288, Novus Biologicals) and mouse Anti-NeuN primary antibody (1:2000, ab104224, Abcam) for one hour at room temperature thereafter. Sections were rinsed three times in PBST subsequently and incubated in a PBST solution containing 3% skim milk powder, 2% normal donkey serum, donkey anti-rabbit IgG Alexa Flour 647 secondary antibody (1:500, Thermo Scientific^TM^) and donkey anti-Mouse IgG Alexa Flour 594 secondary antibody (1:500, Thermo Scientific^TM^) for one hour. Sections were rinsed three times in PBST and one time in PBS before they were mounted on microscope slides, cover slipped with a mounting media containing DAPI (ProLong™ Diamond Antifade Mountant, Thermo Scientific^TM^) and stored at 4°C until imaging under a FluoView FV10i confocal laser scanning microscope (Evident Corporation, Tokyo, Japan). Subjects with missing, incomplete or off-region GFP signaling were excluded from the analyses.

#### Western Blot

Brain punches were lysed in NP40 lysis buffer containing 1% Nonidet P40, 0.1% SDS, 50 mM Tris base, 0.1 mM EDTA, 0.1 mM EGTA, and 0.1% deoxycholic acid, pH 7.4. A cocktail of inhibitors composed of 1 mM sodium pyrophosphate, 1 mM sodium orthovanadate, 20 mM NaF, a protease inhibitor, and a phosphatase inhibitor (Sigma-Aldrich, P5726-1ML) at 1 mM was added to the NP40 buffer. The lysate was incubated under rotation for 30 minutes at 4°C. Proteins were found in the supernatant obtained after centrifugation at 14,000 rpm for 10 minutes at 4°C. Protein quantification was performed using the Pierce BCA Protein Assay Kit (ThermoScientific, #23227) with a BSA standard curve from 0 to 1 μg/μL to obtain a protein concentration of 30 μg. Proteins were mixed with 4X Laemmli loading buffer (Bio-Rad, #1610747) in a total volume of 60 μL and heated to 95°C for 5 minutes to denature the proteins. Protein samples were separated by SDS-PAGE on a 10% acrylamide gel using the Mini-PROTEAN Tetra System (Bio-Rad). Migration was performed in 1X Tris-Glycine electrophoresis buffer for 30 minutes at 100 V to gather the proteins, then one hour at 120 V for migration in the separation gel. The transfer was done onto a 0.2 μm nitrocellulose membrane (Bio-Rad, #1620112) for one hour at 100V in cold 1X Tris-Glycine-methanol transfer buffer.

Membranes were blocked for 1 hour at room temperature in 5% TBST-BSA according to the manufacturer’s recommendations. Blocked membranes were then incubated at 4°C with the primary antibody (2.5% TBST-BSA) until the next day before being incubated with the secondary antibody (1/3000) of rabbit or mouse (Goat Anti-Rabbit IgG (H+L) HRP Conjugate, Millipore, LV1646281, Goat Anti-Mouse IgG (H+L) HRP Conjugate, Millipore, AP124P) for one hour at room temperature. Proteins were detected using the Clarity Western ECL Substrate detection system (Bio-Rad, #170-5061). Primary antibodies used are: Bmal1 (Novus Biologicals, NB100-2288, Dilution 1:1000, Rabbit) and b-actine (Sigma, A5316, Dilution 1 : 2000, Mouse). Densitometric analyses of immunoblots were performed using ImageJ software (Fiji, Version 2.1.0/1.53c).

#### Data analysis and Statistics

Prism 10 (Software, San Diego, CA, USA) was used to statistically analyze and visualize the results of this study. The outcomes were depicted as mean ± standard error of the mean (SEM). Results of behavioural tests were compared by two-way ANOVA (factors: sex and viral vector) followed by Sidak’s multiple comparison. Outcomes of experiments conducted in constant light were compared using a three-way ANOVA (factors: sex, light condition and test phase) and analysed using Sidak’s multiple comparison thereafter. To determine whether time of day affected diurnal levels of striatal DA, a one-way ANOVA was performed. Subsequently, a cosine analysis and zero-amplitude test were conducted to further assess the significance of daily oscillations in DA levels (Cornelissen, 2014). Daily profiles of gene expression and DA levels were analyzed using a two-way ANOVA (factors: viral vector and time point) followed by Tukey’s multiple comparisons if data were normally distributed. The level of significance was set at p ≤ 0.05.

## Author contributions

K.S., S.A., and C.G. conceived and designed the study; C.G., N.B., A.S.., and H.J. performed the experiments; C.G., K.S., and S.A. analyzed and interpreted the data, and wrote the manuscript; S.A. and K.S. Supervision.

## Acknowledgements

This work was funded by grants from the Canadian Institutes of Health Research (S.A). All microscopy was performed at the Concordia Centre for Microscopy and Cellular Imaging at Concordia University, Montreal (special thanks to Dr. Chris Law).

---

**Supplemental Figure 1.**
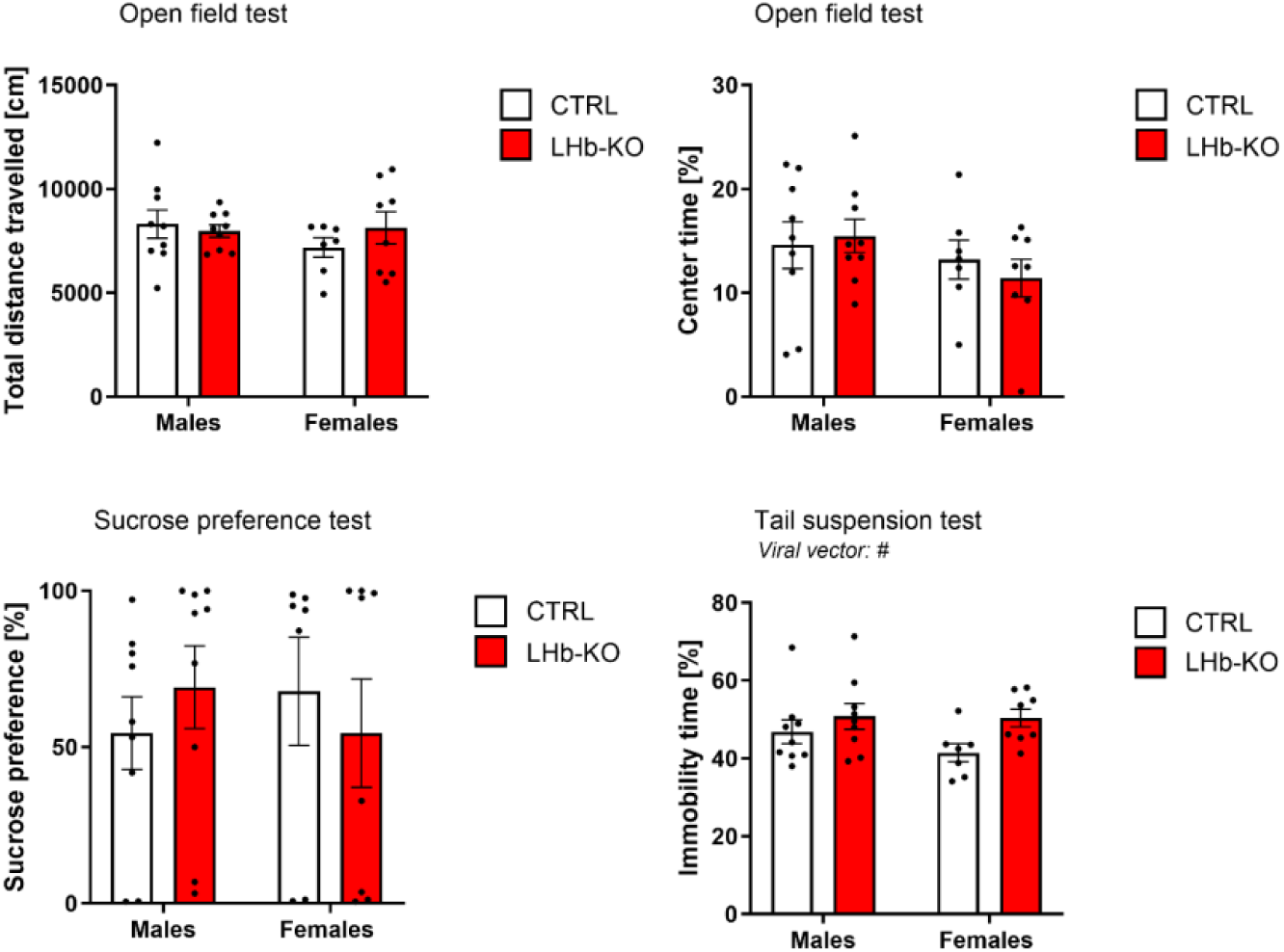
LHb Bmal1 KO does not meaningfully impact affective behaviours. Males: n = 18 (9 KO); females: n = 15 (7 KO). Two-way ANOVAs for Viral vector x sex was run for all tests. A) Open field test, distance travelled. B) Open field tests, time spent in center of the field. C) Sucrose preference test for a 1% sucrose solution. D) Time immobile during the tail suspension test.

